# TileBac: A Benchmark CryoEM Dataset of Bacteria in Ultralow-Dose Montage Tiles

**DOI:** 10.64898/2026.06.08.731030

**Authors:** Lynnicia N. Massenburg, Sita S. Madugula, Spenser R. Brown, Amber N. Bible, Chanda R. Harris, Scott T. Retterer, Jennifer L. Morrell-Falvey, Rama K. Vasudevan, Alexis N. Williams

## Abstract

Current segmentation models are capable of routine identification of biological features in noisy cryogenic electron microscopy (cryoEM) images. However, there are still challenges with complete segmentation of high boundary, thin objects such as bacterial cell envelopes and flagella. Moreover, ultralow-dose cryoEM images pose as an additional challenge to boundary distinctions between the object and background. Here, we present TileBac, a benchmark dataset of ultralow-dose montage tiles of *Pantoea* sp. YR343 to segment bacterial inner and outer membranes for evaluation of model effectiveness. We show that foundation models outperform convolutional neural networks at continuous bacterial cell envelope segmentation despite having lower performance metrics. We release the TileBac benchmark dataset on Hugging Face for further insights into model architecture development.

## 1 Introduction

The cell-surface interface usually involves high-boundary cellular features of interest such as membranes and flagella. Managing the spread of pathogenic bacterial cells across surfaces has been a key factor in biotic-abiotic interface research [1, 2]. Therefore, it is pertinent to understand the morphology of these features as targets for mitigating cell spread interest with a microscopic-scale investigation in cell-surface interactions. Gram-negative bacteria are of particular interest due to the cell envelope sensitivity to environmental stress [3, 4]. We selected *Pantoea* sp. YR343, a Poplar soil microbe [5], as an ideal model to test high boundary, thin objects such as bacterial cell envelopes and flagella in ultralow-dose cryoEM images.

Artificial intelligence-based tools have been developed for bacterial ultrastructure characterization such as bacterial cell envelope thickness, bacteria-flagella interactions and bacteria in a field of view [6]. These tools were built with low-dose cryoEM images at 40 e^−^/Å^2^ and used an input of segmentation masks generated by a fine-tuned pre-trained YOLOv11 segmentation model. Ongoing work in progress has since shown that low-dose cryoEM images are compatible with artificial intelligence (AI) model architectures, but lower electron dose imaging conditions may reduce model performance [7].

Stitched montage images are ideal for capturing high-resolution information in a wide field of view to see biofilm interactions on surfaces. However, this imaging routine has the caveat of ultralow-dose imaging below 10 e^−^/Å^2^ to maintain efficient speeds in a tile set collection. This dataset contains ultralow-dose (4 e^−^/Å^2^) cryoEM montage tile image and SAM3 segmentation models have been fine-tuned to predict bacterial inner membranes and outer membranes. This bacterial membrane dataset called TileBac is a benchmark dataset to challenge current AI workflows in rapid, seamless montage stitching and stitched segmentation of high-boundary objects in extremely noisy ultralow-dose cryoEM images. Bacterial flagella low-dose cryoEM movies (40 e^−^/Å^2^ total dose with 40 fractions at 1 e^−^/Å^2^ per fraction) have also been added as a dataset challenge to segmenting high-boundary thin objects in user-averaged ultralow-dose cryoEM images.

## 2 Related Work

### 2.1 Microbial CryoEM Datasets

Object detection and semantic segmentation in electron microscopy (EM) offers strategies to auto-label biological features in EM images to reduce the manual annotation time costs to use for model training [8, 9]. Current strategies use zero-shot Segment Anything Model (SAM) model architecture from Meta AI for object detection and segmentation in SAM-EM, MicroSAM and CellSAM for liquid phase electron microscopy [10–12]. U-Net-like convolutional neural networks are also well-established to segment eukaryotic cell features [13, 14]. Availablity of bacteria-specific cryoEM datasets remain limited. Few are stored on EMPIAR as a public archive. This dataset serves as an untapped resource for training on bacterial membrane features [15]. For montage-type datasets, volume electron microscopy routinely tiles and stitches images for large 2-dimensional and 3-dimensional montage views [16]. YOLO-based stitched segmentation with SAHI are now conventional workflows for tiling and stitching for object detection [17–19]. Ultralow-dose bacterial images provided in this work represents a rare dataset release for endeavours in image registration (seamless tiling) and stitched segmentation in ultralow-dose cryoEM images of bacteria.

## 3 Dataset

This dataset represents the first benchmark dataset to combine ultralow-dose imaging with image tiling to benchmark thin biological object segmentation. Here we showcase TileBac, a collection of cryoEM images from a benchmark montage tile dataset of *Pantoea* sp. YR343, a soil microbe [5]. Figure 1 shows representative images of the dynamic normalized images at 4096 × 4096 at 4 e^−^/Å^2^ total electron dose per image. These images were tiled to 2048 × 2048 to increase the pixel coverage of the membrane boundary during bilinear resizing in model training and inference. The image tiling also reduced the background presence as tiled background-only images were removed from the training set to prevent background oversaturation. This dataset focuses on the inner membrane and outer membrane as two thin-object classes of interest.

**Figure 1:**
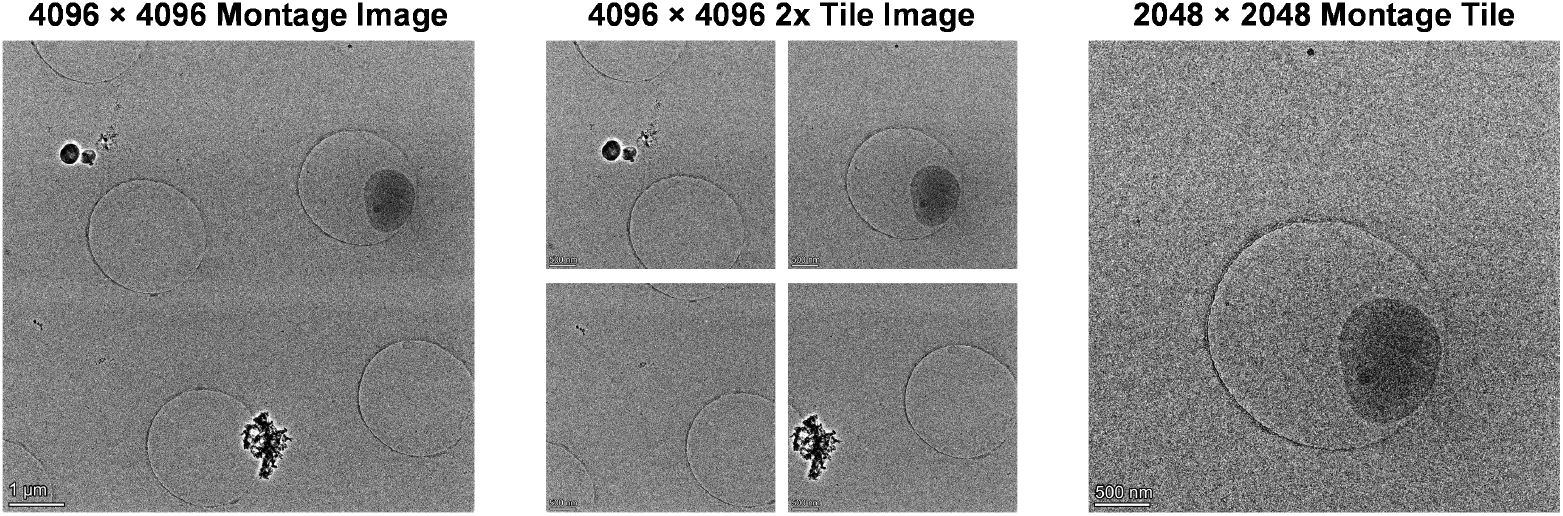
Representative ultralow-dose cryoEM montage images and tiles TileBac benchmark dataset. Example post-processed ultralow-dose 4096 × 4096 montage image is shown in the left panel; scale bar is 1 *µ*m. The center panel shows further tiling of the montage image to 2048 × 2048 montage tiles to reduce thin membrane pixel compression during image downsizing; scale bar is 500 nm. Example cryoEM image in the TileBac benchmark dataset.

### 3.1 Data collection

TileBac is designed to benchmark segmentation, detection, and robustness of AI models under ultralow-dose cryoEM imaging conditions. Below we describe the dataset collection details for ultralow-dose imaging conditions in this dataset. *Pantoea* sp. YR343 were grown in R2A or MOPS + glucose before Vitrobot vitrification. For each condition, 3 *µ*L aliquots were applied to glow-discharged Quantifoil® or C-Flat 2/2 gold grids, blotted using a Vitrobot Mark IV (Thermo Fisher Scientific) at a blot force of −5 for 5–10 s at 100% humidity and 4.5 °C, and vitrified by plunge-freezing in liquid ethane/propane cooled with liquid nitrogen. Montage images were automatically collected on Thermo Fisher Scientific Krios G4 in nanoprobe TEM mode with 70 *µ*m C2 aperture under cryogenic conditions. The Falcon 3EC direct electron detector (counting mode) with a total electron dose of 4 e^−^/Å^2^ per montage image. These images are called ultralow-dose montage (ULDM) images. For montage image automation, an automated montage search map was collected using Tomography 5 (Thermo Fisher Scientific) 7 × 7 montage tiles at 4096 × 4096 image size (2.128 nm/pix). These parameters were chosen to cover the space of a grid square for a 200-mesh size grid.

Stitched montages were automatically generated at 12005 × 12005 image size (25 nm/pix) in Tomography 5. Figure 2 shows example montage images before and after dynamic normalization pre-processing used to enhance image contrast and image stitching to generate a complete stitched montage image. However, the bacterial cell size is reduced in the stitched montage image compared to the original montage image (Figure 2). The reduced size of the bacterial membranes in the stitched montage image will pose a significant deterrent to membrane identification, hence the stitched montage images were not used as the benchmark dataset. Images were collected as .mrc files and converted to .png files for dynamic normalization prior to tiling to 2048 × 2048 tiles (Figure 1 and Figure 2). To further increase the bacterial cell size with respect to model-specific image downsizing, the original montage images were further tiled from 4096 × 4096 to 2048 × 2048 montage tiles. Hence, our benchmark dataset is constructed using 2048 × 2048 tiles from the 4096 × 4096 montage images.

**Figure 2:**
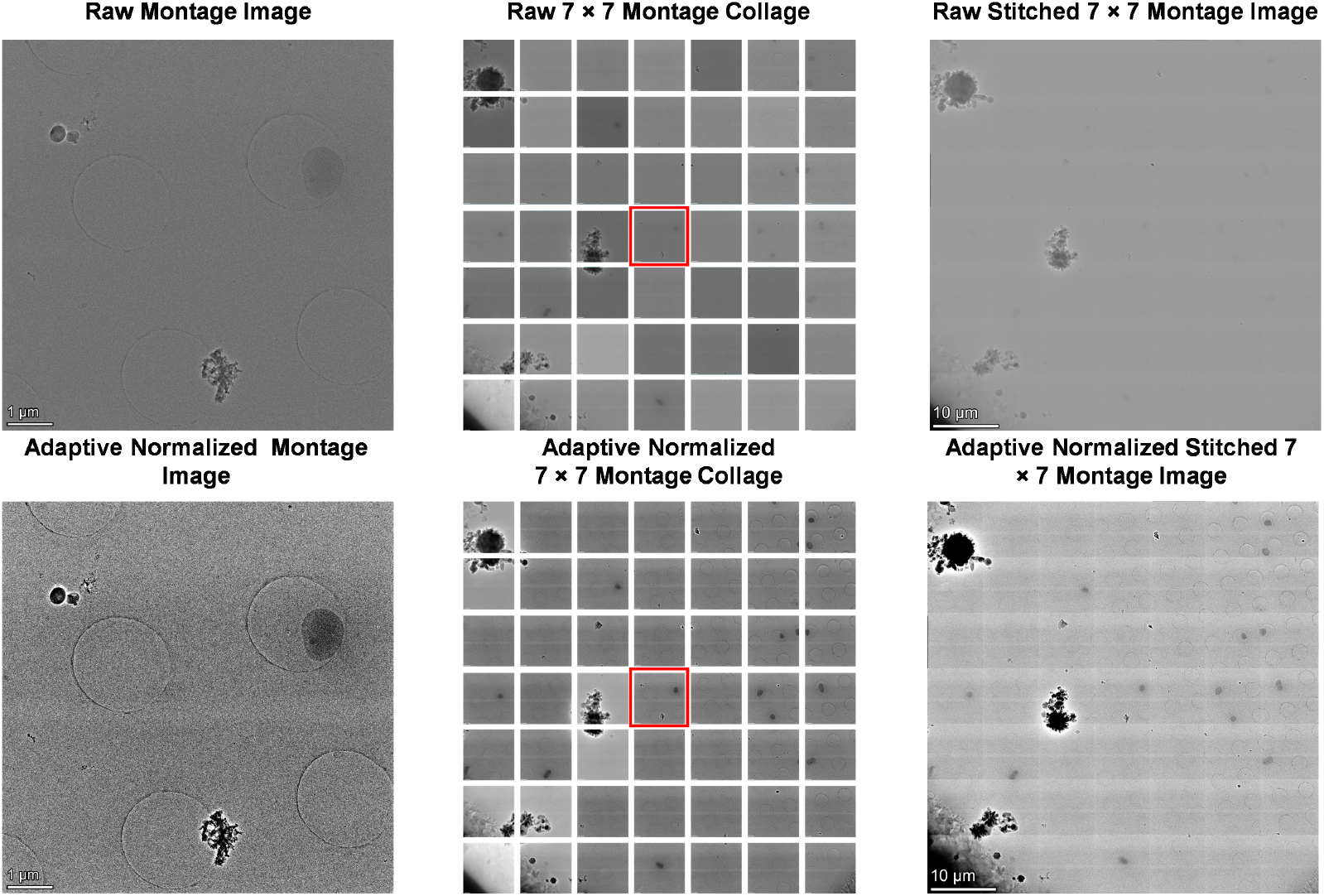
Dynamic normalization in montage tiles and montage-stitched images in EPU. Left panel shows 4096 × 4096 montage tile before (top) and after (bottom) adaptive (or dynamic) normalization. The middle panel shows the montage image collage before image stitching. The image in the red box represents the montage tile in the left panel. The stitched montage tile (12005 × 12005) automatically generated from the Tomography 5 Search Map workflow.

### 3.2 Data organization and annotation

The TileBac dataset is manually annotated in Roboflow and organized in an 80:5:15 train, valid and test split (Figure 3) [20]. The validation set is kept smaller than the held-out test set to check for robust unbiased model benchmarking as an initial challenge on model robustness for 2048 × 2048 ultralow-dose image montage tiles. Annotated inner membranes (IM) are labelled class 0 and outer membranes (OM) are class 1. All images are resized to either 640 × 640 or 1024 × 1024 to track model performance in bilinear interpolation. Augmentations and resizing of the training images were performed as the presence of bacteria was sparse in our montage tile sets (Table 1). Annotated images are output in either YOLO text format for YOLOv11 and YOLO26 segmentation models or COCO format for U-Net, Detectron2 and SAM3 segmentation models.

**Table 1:**
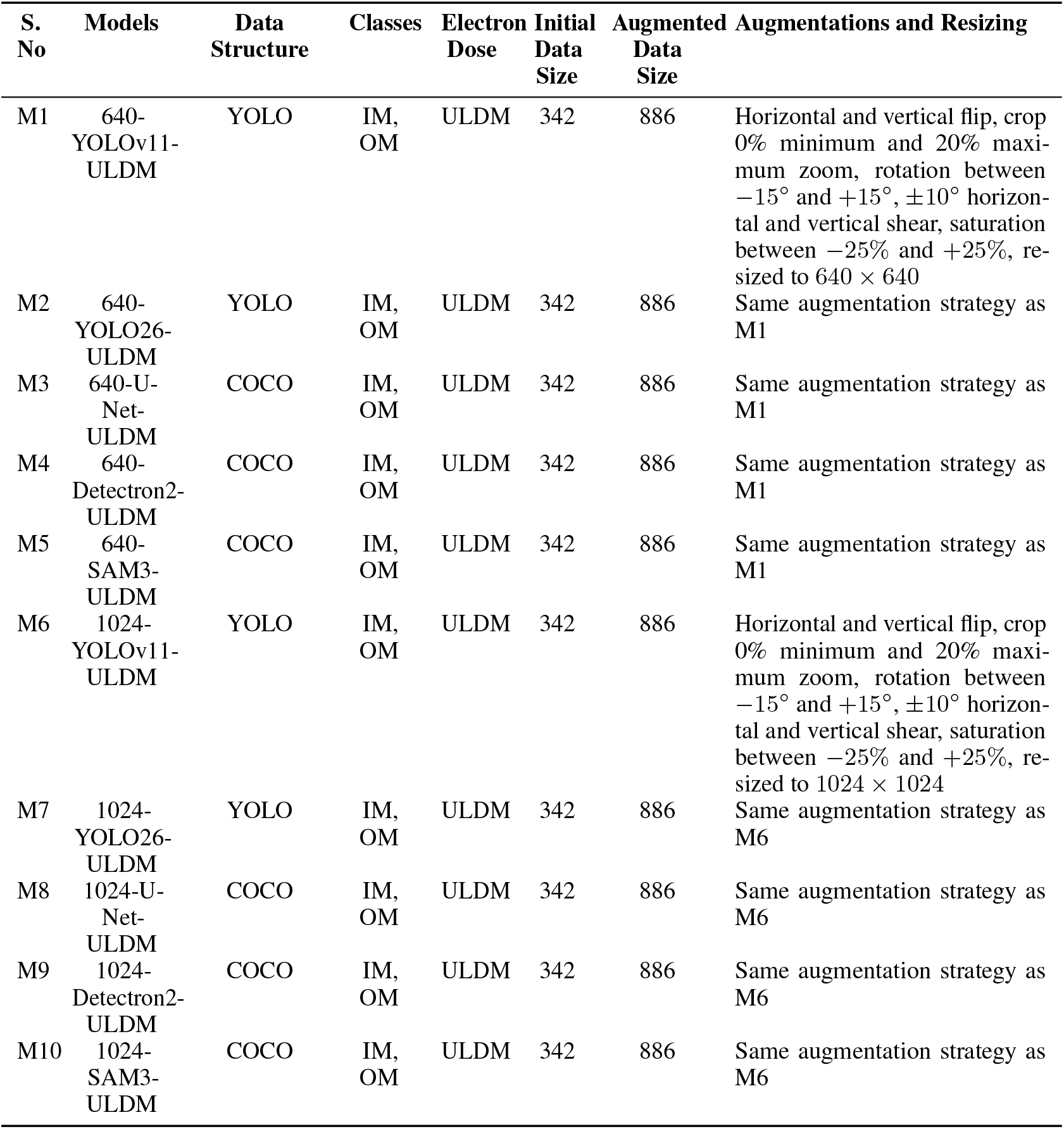
Summary for the ULDM tile dataset, augmentations, and resizing applied for segmentation model training of the outer membrane and inner membranes.

**Figure 3:**
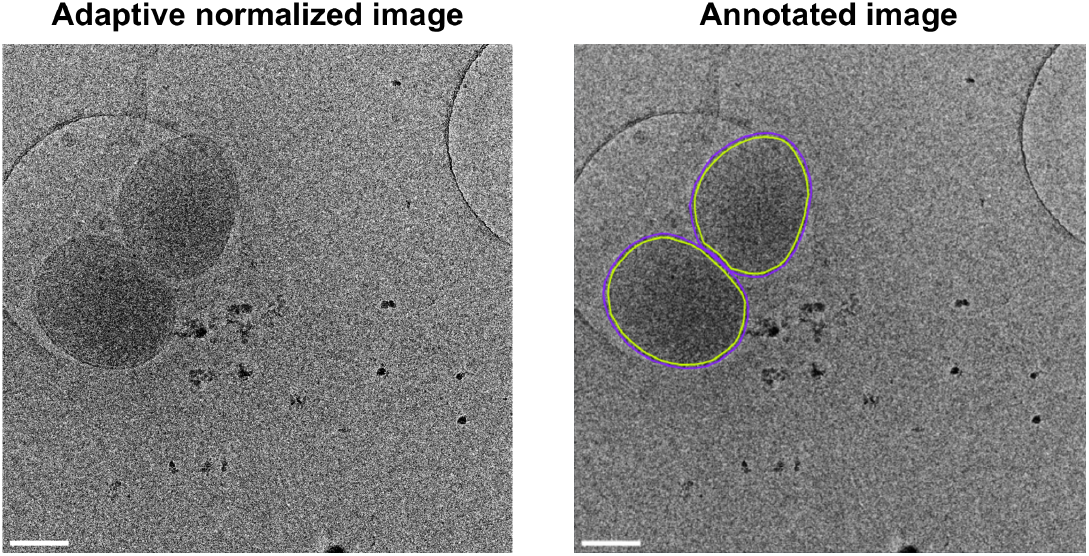
Annotation of bacteria inner and outer membranes in ultralow-dose montage tiles. Montage tiles (2048 × 2048, left panel) are annotated based on bacteria inner membranes (lime) or outer membranes (purple, right panel). Scale bar is 500 nm.

## 4 Experiment

### 4.1 Implementation details

A collection of segmentation model architectures were used for the identification of bacterial membranes: YOLOv11, YOLO26, U-Net, Detectron2 and SAM3[17, 18, 13, 21–24]. Google Colab (A100 GPU) was used to fine-tune pre-trained models. Table 2 outlines the training hyperparameters used to benchmark segmentation models.

**Table 2:** Training hyperparameters for benchmarked segmentation models.

| Model | Optimizer | Learning Rate (LR) | Batch Size | Epochs / Iterations | Input Size | Selection Criterion |
| --- | --- | --- | --- | --- | --- | --- |
| YOLOv11n-seg | AdamW | $1 \times 10^{-3}$ | 16 | 100 epochs | 640 / 1024 | Best mAP50–95 |
| YOLO26n-seg | AdamW | $1 \times 10^{-3}$ | 16 | 100 epochs | 640 / 1024 | Best mAP50–95 |
| U-Net | Adam | $1 \times 10^{-4}$ | 4 | 180 epochs | 640 / 1024 | Lowest Dice-BCE loss |
| Detectron2 | Stochastic Gradient Descent | $2.5 \times 10^{-4} \rightarrow 1 \times 10^{-4}$ | 2 | 8,000 iterations | 640 / 1024 | Highest AP |
| SAM3 | AdamW | $5 \times 10^{-7}$ (global), $2.5 \times 10^{-5}$ backbone | 4 | 20 epochs | 1008 | Lowest validation Dice |

### 4.2 Evaluation

Model performance is evaluated on valid and test images using mAP50 for segmentation models and the best F1-score (mathematically equivalent to Dice Coefficient, reported F1-score for classification-based evaluation consistency) for all segmentation models (Table 3 and Table A1). We use the following equation to calculate mAP50:

**Table 3:** Performance metrics on the valid images for the 2048 *×* 2048 ULDM tile fine-tuned models.

| Model | Electron Dose | Class | Mask Precision* | Mask Recall* | mAP50* | Best F1-Score* |
| --- | --- | --- | --- | --- | --- | --- |
| 640-YOLOv11-ULDM | ULDM | All | 0.997 | 1.000 | 0.995 | 0.998 |
|  |  | IM | 0.996 | 1.000 | 0.995 | 0.998 |
|  |  | OM | 0.997 | 1.000 | 0.995 | 0.999 |
| 1024-YOLOv11-ULDM | ULDM | All | 0.997 | 1.000 | 0.995 | 0.999 |
|  |  | IM | 0.997 | 1.000 | 0.995 | 0.999 |
|  |  | OM | 0.997 | 1.000 | 0.995 | 0.998 |
| 640-YOLO26-ULDM | ULDM | All | 0.988 | 0.959 | 0.994 | 0.973 |
|  |  | IM | 0.976 | 1.000 | 0.995 | 0.988 |
|  |  | OM | 1.000 | 0.917 | 0.993 | 0.957 |
| 1024-YOLO26-ULDM | ULDM | All | 0.992 | 1.000 | 0.995 | 0.996 |
|  |  | IM | 0.989 | 1.000 | 0.995 | 0.994 |
|  |  | OM | 0.989 | 1.000 | 0.995 | 0.998 |
| 640-U-Net-ULDM | ULDM | All | 0.987 | 0.988 | – | 0.987 |
|  |  | IM | 0.986 | 0.987 | – | 0.986 |
|  |  | OM | 0.987 | 0.990 | – | 0.988 |
| 1024-U-Net-ULDM | ULDM | All | 0.984 | 0.988 | – | 0.986 |
|  |  | IM | 0.980 | 0.987 | – | 0.984 |
|  |  | OM | 0.987 | 0.989 | – | 0.988 |
| 640-Detectron2-ULDM | ULDM | All | 0.906 | 0.942 | 0.948 | 0.924 |
|  |  | IM | 0.821 | 0.900 | 0.860 | 0.859 |
|  |  | OM | 0.990 | 0.979 | 0.974 | 0.984 |
| 1024-Detectron2-ULDM | ULDM | All | 0.521 | 0.729 | 0.607 | 0.608 |
|  |  | IM | 0.379 | 0.760 | 0.531 | 0.506 |
|  |  | OM | 0.674 | 0.881 | 0.705 | 0.764 |
| 640-SAM3-ULDM | ULDM | All | 0.995 | 0.988 | 1.000 | 0.991 |
|  |  | IM | 0.993 | 0.986 | 1.000 | 0.990 |
|  |  | OM | 0.995 | 0.990 | 1.000 | 0.993 |
| 1024-SAM3-ULDM | ULDM | All | 0.995 | 0.984 | 1.000 | 0.989 |
|  |  | IM | 0.996 | 0.976 | 1.000 | 0.986 |
|  |  | OM | 0.994 | 0.991 | 1.000 | 0.992 |

\*All instance-based model outputs were calculated at IoU@0.5 with a swept confidence threshold during instance-level evaluation for mAP50. All metrics are globally pooled.

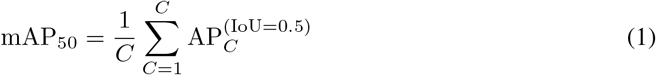

where C is the number of classes, and AP is the area under the Precision-Recall (PR) curve in which AP is described as:

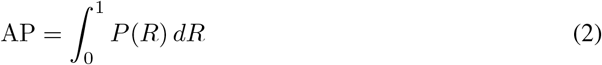

where P is precision, and R is recall. Precision and recall are described as:

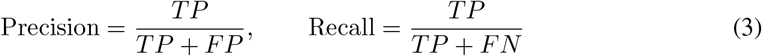

where TP is true positive, FP is false positive and FN is false negative. The F1-score (F1) is therefore described as:

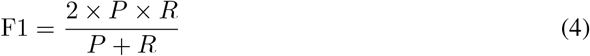

The YOLOv11 architecture showed the highest F1-scores for valid images in all classes, yet showed discontinuous outer membrane masks. However, the test image overall metrics underperformed due to low OM metrics with OM mAP50 scores below 0.65. SAM3 architecture had lower F1-scores, but consistently higher mAP50 scores and continuous filled masks in both valid and test image sets. U-Net showed inconsistencies in mask generation such as incomplete masks and object merging for bacterial neighbours. Detectron2 showed more consistent mask inferences for models trained and inferenced at 640 image size, and inconsistent masks at 1024 image size (Figure 4). See Appendix A (Figure A1 and Figure A2) for PR curve and F1-score curve performance metrics for valid and test image sets.

**Figure 4:**
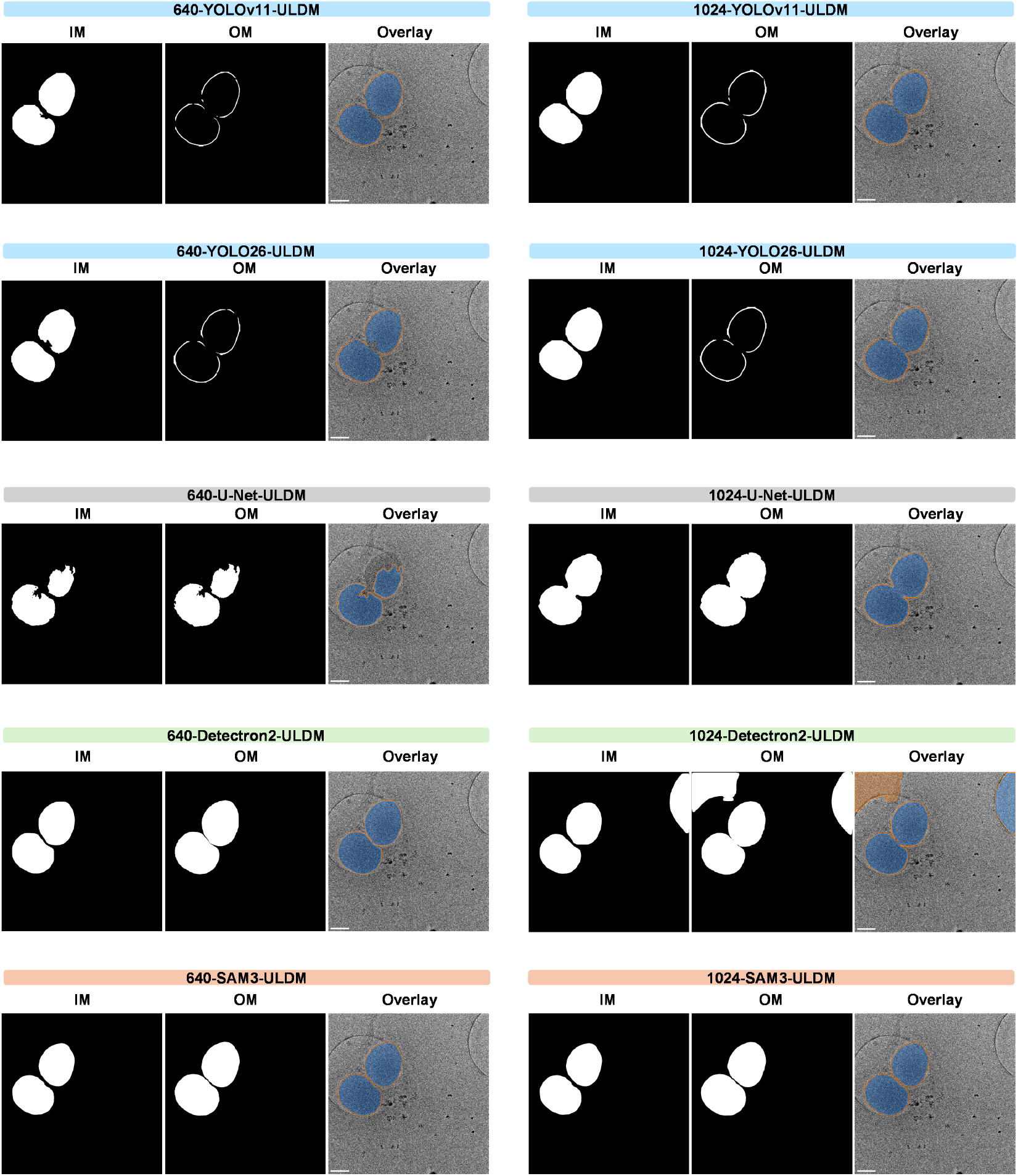
Prediction of bacteria inner and outer membranes in ultralow-dose montage tiles. YOLOv11, YOLO26, U-Net, Detectron2 and SAM3 pre-trained models were fine-tuned with train images with either 640 × 640 or 1024 × 1024 image size. Inferences were output as binary masks and color overlay of bacteria inner membranes (IM, blue) or outer membranes (OM, orange).

#### 4.2.1 Challenge Task 1: Unified model for complete segmentation with 4096 × 4096 montage images to identify bacterial membranes

This challenge will focus on complete mask generation of bacterial membranes with a unified segmentation model. We present benchmarks for YOLOv11, YOLO26, U-Net, Detectron2 and SAM3 models to showcase model performance in a range of architectures. Overall, these models showed mask imperfections even with high performing metrics with valid images. Metrics underperformed in test images with the exception of the SAM3 foundation model. We have not applied this workflow to the original 4096 × 4096 montage images as a challenge to the presented model architectures. Several bypasses to this challenge include tiling to 2048 × 2048 performed in this work, stitched segmentation or sliding window segmentation of the 4096 × 4096 image to enlarge objects of interest for model performance improvement. However, these approaches can be time consuming to process multiple images and recompile the predicted masks on the original image. Foundational models may be ideal in this ultralow-dose cryoEM dataset, yet may not show consistency across heterogenous EM datasets [25]. We seek a unified model with a convolutional neural network architecture focused on model improvement of thin membrane objects on the original 4096 × 4096 montage images with a streamlined inference pipeline for speed and compute efficiency.

#### 4.2.2 Challenge Task 2: Create an image processing pipeline that seamlessly registers the montage image collage that does not cause overlap issues with bacterial membranes, then tiles the stitched montage collage, inferences the tiles and segmentation-stitches the predicted masks to identify bacterial membranes

We seek to focus on stitched segmentation of bacterial membranes on this challenge to overcome large image sizes with stitched montage images. Current montage registration routines cause slight overlap issues that misalign the bacterial membrane, posing as a challenge in bacterial membrane identification currently being addressed in literature [26]. After image registration the montage array into a stitched montage, the resulting image size could be as large as 12005 × 12005 pixel dimensions that can cause detrimental image compression and loss of features of interest in model inferencing. Therefore, re-tiling is often needed before resizing to 640 × 640 or 1024 × 1024. Create a workflow for seamless montage stitching with a field of view, re-tiling during inference and stitched segmentations with a model robust to noisy cryoEM images.

#### 4.2.3 Challenge Task 3: Create a unified model that is robust to montage image scale variation to identify the bacterial membranes

The third challenge is focused on creating a model with the flexibility of segmenting bacterial membranes at different image scales, a combination and extension of Challenge Task 1 and 2.

Specifically, the model must accurately segment both large bacterial structures in montage images and small bacterial structures in stitched collage images. Montage imaging enables higher-resolution visualization across a field of view on a grid square compared to a single grid square image. Key features of interest include the bacterial inner membrane and outer membrane that are more clearly resolved at the montage image scale. Challenge Task 1 focuses on these large-scale bacterial structures. In standard montage stitching workflows in Tomography 5, montage images are arranged as tiles and stitched to a single montage collage image. This image is often re-tiled to meet model input constraints as described in Challenge Task 2 resulting in smaller object sizes than in Challenge Task 1. Consequently, models trained for large-object segmentation may not generalize to smaller-scale representations, often requiring separate models for each scale.

This challenge therefore proposes the development of a unified model that is robust to scale variation, enabling accurate segmentation of both large bacterial objects in montage images and small bacterial objects in stitched montage collage images. Several convolutional neural networks have shown promise in multi-scale image segmentation [27–29]. This task is particularly difficult because the bacterial cell membrane is inherently a thin structure, making this feature highly sensitive to scale-dependent resolution changes. An all-scale model will expand use for not only individual bacteria, but also large spans of bacterial clusters.

## 5 Conclusions

### 5.1 Summary and limitations

This paper presents TileBac, a benchmark dataset used to determine the segmentation model performance in noisy ultralow-dose cryoEM images to segment bacterial inner and outer membranes. We also included a second flagella dataset for benchmarking high-boundary flagellar objects in user-averaged ultralow-dose conditions. For limitations, our benchmark dataset is only relevant for Gram negative bacteria. In addition, the presence of partial cells due to montage tile boundaries may inadvertently lower model benchmarking performance. For ethical considerations, this work may be used to image pathogenic bacteria for segmentation and feature discovery. Even though *Pantoea* is a plant soil microbe, strains of *Pantoea* have been associated with diseases in humans [30]. This dataset will be applicable to study both beneficial and pathogenic bacteria, and care must be taken with future data analysis to mitigate bio-weapon characterization and to advance the medical safety of the society at large. Overall, this dataset is a versatile testing ground for the development of a high-precision model for thin object segmentation in low-contrast cryoEM images.

### 5.2 Broad impacts

Medical interventions can be designed around the reduction of biofilm attachment to high-touch surfaces in medical facilities. Flagella may play a role in surface adhesion of bacteria and biofilm establishment by promoting adhesion to hydrophobic surfaces, a key target in mitigating bacterial attachment to surfaces [31–35]. We provide additional dataset processing challenges with our flagella dataset focused on the segmentation of thin high-boundary flagellar objects in Appendix B.

## Acknowledgments and Disclosure of Funding

This work is supported by the U.S. Department of Energy, Office of Science FWP ERKCZ64, Structure Guided Design of Materials to Optimize the Abiotic-Biotic Material Interface, as part of the Biopreparedness Research Virtual Environment (BRaVE) initiative. Sample preparation, imaging and image analysis were conducted as part of a user project at the Center for Nanophase Materials Sciences (CNMS), which is a US Department of Energy, Office of Science User Facility at Oak Ridge National Laboratory. Electron microscopy data was collected using instrumentation within ORNL’s Materials Characterization Core provided by UT-Battelle, LLC, under Contract No. DE-AC05-00OR22725 with the DOE and sponsored by the Laboratory Directed Research and Development Program of Oak Ridge National Laboratory, managed by UT-Battelle, LLC, for the U.S. Department of Energy.

## Public Disclosure

This manuscript has been authored by UT-Battelle, LLC, under contract DE-AC05-00OR22725 with the US Department of Energy (DOE). The United States Government retains and the publisher, by accepting the article for publication, acknowledges that the United States Government retains a nonexclusive, paid-up, irrevocable, worldwide license to publish or reproduce the published form of this manuscript, or allow others to do so, for the United States Government purposes. The Department of Energy will provide public access to these results of federally sponsored research in accordance with the DOE Public Access Plan(http://energy.gov/downloads/doe-public-access-plan).

## Data Availability Statement

All datasets are openly available on Constellation (DOI: 10.13139/ORNLNCCS/3025229). A description of TileBac can be found on https://github.com/Lynnicia/TileBac-ultralow-dose-montage-tiles. Ultralow-dose montage datasets and raw images can be found on Hugging Face at: https://huggingface.co/datasets/LynnMass/tilebac-ULDM-benchmark-dataset, https://huggingface.co/datasets/LynnMass/tilebac-ULDM-tiles, and https://huggingface.co/datasets/LynnMass/tilebac-stitched-montage. Flagella datasets can be found on Hugging Face at: https://huggingface.co/datasets/LynnMass/tilebac-flag-dataset. Ultralow-dose montage segmentation models can be found at: https://huggingface.co/LynnMass/640-YOLOv11-ULDM, https://huggingface.co/LynnMass/1024-YOLOv11-ULDM, https://huggingface.co/LynnMass/640-YOLO26-ULDM, https://huggingface.co/LynnMass/1024-YOLO26-ULDM, https://huggingface.co/LynnMass/640-U-Net-ULDM, https://huggingface.co/LynnMass/1024-U-Net-ULDM, https://huggingface.co/LynnMass/640-Detectron2-ULDM, https://huggingface.co/LynnMass/1024-Detectron2-ULDM, https://huggingface.co/LynnMass/640-SAM3-ULDM and https://huggingface.co/LynnMass/1024-SAM3-ULDM.

## A Appendix: Metrics Performance for Ultralow-Dose Montage Tiles

**Table A1:** Unbiased performance metrics on the hold-out test images for the 2048 × 2048 ULDM tile fine-tuned models.

| Model | Electron Dose | Class | Mask Precision* | Mask Recall* | mAP50* | Best F1-Score* |
| --- | --- | --- | --- | --- | --- | --- |
| 640-YOLOv11-ULDM | ULDM | All | 0.998 | 1.000 | 0.995 | 0.999 |
|  |  | IM | 0.997 | 1.000 | 0.995 | 0.999 |
|  |  | OM | 0.998 | 1.000 | 0.995 | 0.999 |
| 1024-YOLOv11-ULDM | ULDM | All | 0.997 | 1.000 | 0.995 | 0.999 |
|  |  | IM | 0.997 | 1.000 | 0.995 | 0.999 |
|  |  | OM | 0.997 | 1.000 | 0.995 | 0.998 |
| 640-YOLO26-ULDM | ULDM | All | 0.990 | 0.971 | 0.994 | 0.980 |
|  |  | IM | 0.980 | 1.000 | 0.995 | 0.990 |
|  |  | OM | 1.000 | 0.943 | 0.993 | 0.971 |
| 1024-YOLO26-ULDM | ULDM | All | 0.993 | 1.000 | 0.995 | 0.997 |
|  |  | IM | 0.990 | 1.000 | 0.995 | 0.995 |
|  |  | OM | 0.990 | 1.000 | 0.995 | 0.998 |
| 640-U-Net-ULDM | ULDM | All | 0.988 | 0.989 | – | 0.988 |
|  |  | IM | 0.987 | 0.988 | – | 0.987 |
|  |  | OM | 0.989 | 0.991 | – | 0.990 |
| 1024-U-Net-ULDM | ULDM | All | 0.985 | 0.989 | – | 0.987 |
|  |  | IM | 0.981 | 0.988 | – | 0.985 |
|  |  | OM | 0.988 | 0.990 | – | 0.989 |
| 640-Detectron2-ULDM | ULDM | All | 0.912 | 0.948 | 0.952 | 0.930 |
|  |  | IM | 0.828 | 0.907 | 0.865 | 0.866 |
|  |  | OM | 0.995 | 0.989 | 0.978 | 0.992 |
| 1024-Detectron2-ULDM | ULDM | All | 0.534 | 0.741 | 0.618 | 0.620 |
|  |  | IM | 0.391 | 0.771 | 0.542 | 0.519 |
|  |  | OM | 0.687 | 0.889 | 0.716 | 0.775 |
| 640-SAM3-ULDM | ULDM | All | 0.996 | 0.989 | 1.000 | 0.992 |
|  |  | IM | 0.994 | 0.987 | 1.000 | 0.991 |
|  |  | OM | 0.996 | 0.991 | 1.000 | 0.994 |
| 1024-SAM3-ULDM | ULDM | All | 0.996 | 0.985 | 1.000 | 0.990 |
|  |  | IM | 0.997 | 0.977 | 1.000 | 0.987 |
|  |  | OM | 0.995 | 0.992 | 1.000 | 0.993 |

\*All instance-based model outputs (YOLO, Detectron2, and SAM3) were calculated at IoU@0.5 with a swept confidence threshold during instance-level evaluation for mAP50. All metrics are globally pooled.

**Figure A1:**
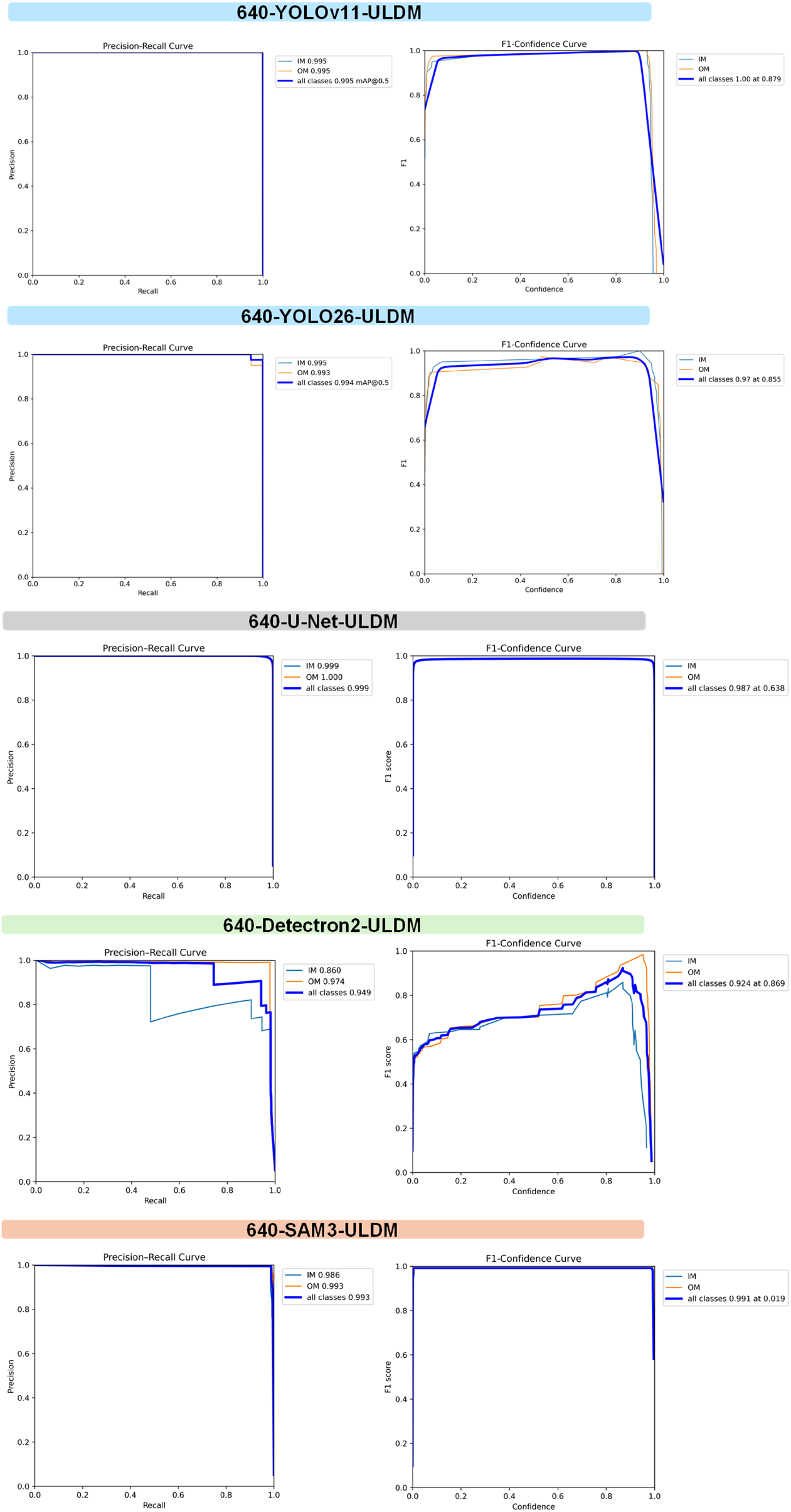
Performance metrics for YOLOv11, YOLO26, U-Net, Detectron2 and SAM3 trained on resized 640 × 640 ULDM tiles.

**Figure A2:**
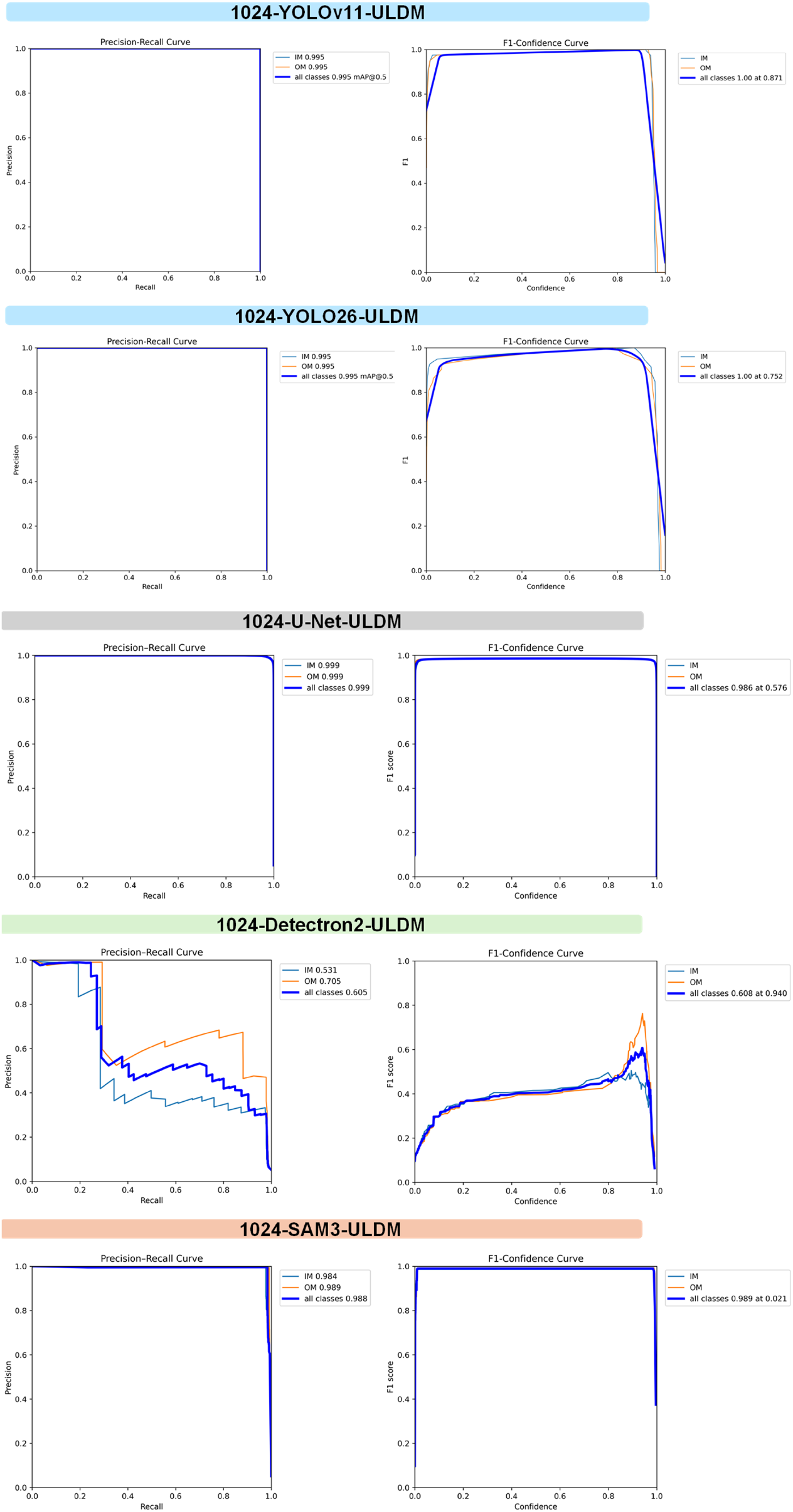
Performance metrics for YOLOv11, YOLO26, U-Net, Detectron2 and SAM3 trained on resized 1024 × 1024 ULDM tiles.

## B Appendix: Low-Dose and Ultralow-Dose Bacteria and Flagella CryoEM Images

### B.1 Low-Dose Dataset: Bacteria and Flagella CryoEM Images

This flagella dataset includes cryoEM images of *Pantoea* sp. YR343 bacteria images manually screened for the presence of flagella. Flagella dimensions typically have a diameter of 12-30 nm and a length beyond 5 *µ*m [36]. *Pantoea* typically have one polar flagella attached with possible secondary polar extensions. However, from our dataset, it is difficult to discern if the flagella are, in fact, attached to the bacteria or overlapping with the bacteria. For clarity, we present this dataset as manually collected cryoEM images of flagella overlapping with bacteria. Similar cryoEM imaging and segmentation workflows have previously been applied to segment bacterial outer membranes and flagella and quantitatively characterize bacteria and flagella overlap [6].

#### B.1.1 Data Collection and Annotation

The annotated cryoEM images used in this dataset are not from an automated montage dataset as is the focus of this paper, but rather a manual collection of bacteria and flagella in low-dose cryoEM. Due to the length of the flagella described above, flagella are an ideal sample for a future montage tile collection. *Pantoea* sp. YR343 were grown in R2A, MOPS + glucose or MOPS + succinate before Vitrobot vitrification as described in Section 3.1. These manually collected flagella were imaged at 40 e^−^/Å^2^ at 0.8986 nm/pix (for 4096 × 4096 images). These images were collected on the Thermo Fisher Scientific Krios G4 with a Falcon 3EC (Thermo Fisher Scientific, counting mode) and cryogenic temperatures. Images were pre-processed by converting .mrc files to .png files and enhancing image contrast with dynamic normalization (per-image adaptive normalization), followed by manual annotation of bacterial cells (outer membrane annotation only) and flagella in Roboflow (Figure B1) [20]. Low-dose adaptive normalization has less noticeable contrast enhancements compared to the noisy ultralow-dose montage tiles in Figure 2. Data are annotated as flagella (class 0) and bacteria (class 1) with an 80:10:10 train, valid and test split. Additional image outputs are summarized in Table B1 describing the resizing and augmentations. Both 2x-train and 3x-train outputs were generated to overcome the low frequency of flagella present in the dataset.

**Figure B1:**
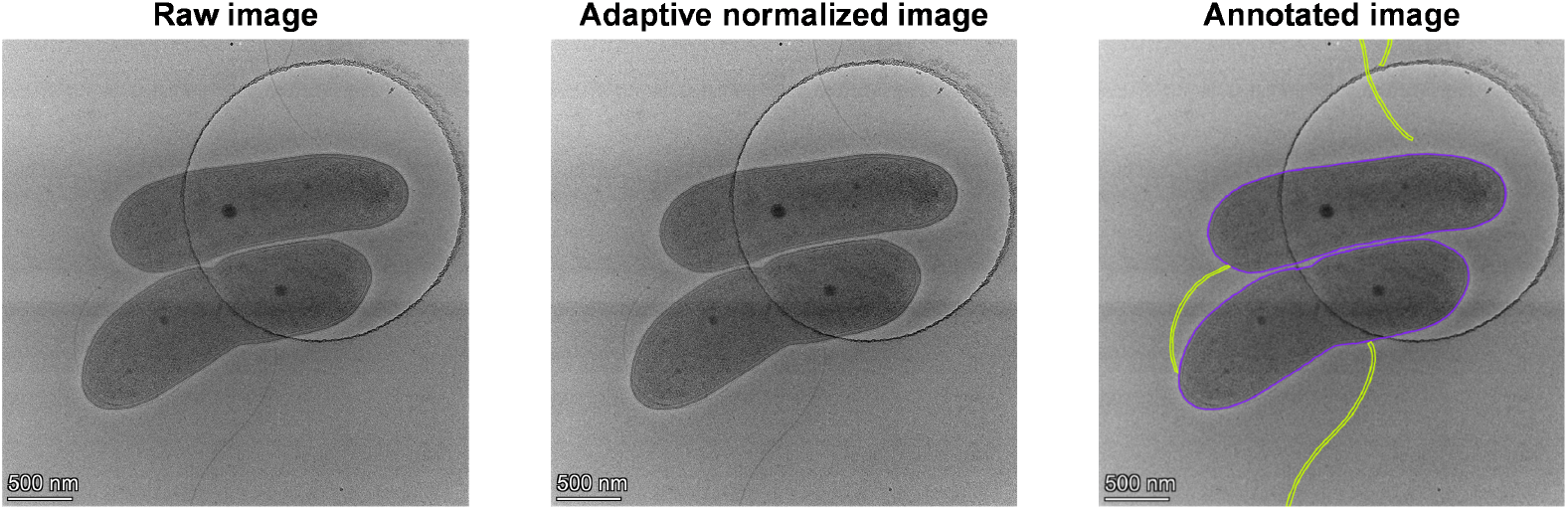
Representative low-dose cryoEM images of bacteria and flagella. Raw (left panel), adaptive normalized (center panel) and annotated (right panel) images of bacteria (purple outline) and flagella (lime outline).

#### B.1.2 Challenge Task B1: Unified Model for Continuous Flagella Segmentation in Low-Dose Conditions

A previous study has trained segmentation models with a bacterial dataset to detect bacteria and flagella [6]. We evaluated the models on our imaging conditions and observed that there are cases of discontinuous flagella segmentation even though the model successfully contoured the flagella (Figure B2). We imaged a low-dose cryoEM dataset consisting of bacteria and flagella as a representative dataset to screen for thin object performance with segmentation models. Given the discontinuous segmentation in the previously published model, we concluded that a unified model is needed to address noisy cryoEM images. We state the same task with bacterial flagella to identify a new model architecture that can be boundary-aware of the entire presented flagella.

**Figure B2:**
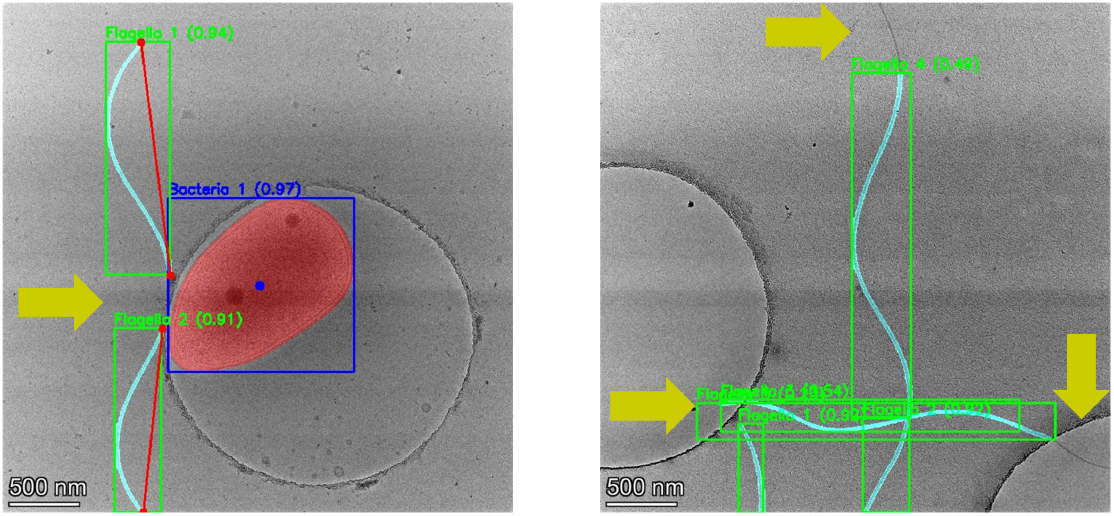
Representative images of bacteria and flagella segmentation. The left panel shows high flagella confidence with YOLOv11 object detection, but segmented flagella is prone to discontinuous segmentation (yellow arrow). The right panel shows incomplete flagella detection (yellow arrows) and redundant prediction of the same flagellum.

### B.2 Ultralow-Dose Dataset: Bacteria and Flagella CryoEM Movies

Due to the length of flagella spanning over 5 *µ*m, montage tiling is an ideal solution to capture the full length of the flagella. However, as described in the main paper, montage tiling often involves ultralow-dose modes for efficient speed in automated tiling data collection. We created a dataset to customize an ultralow-dose mode investigation to identify bacteria and flagella.

#### B.2.1 Data Collection

We collected the same flagella datasets as 40-frame movies (.mrc files) at 1 e^−^/Å^2^ frame in counting mode on the Falcon 3EC. This frame partitioning allows for a user to output averaged images between a range of 1 and 40 e^−^/Å^2^ total electron dose (Figure B3). As such, these raw frames can be grouped at four frames for a total of 4 e^−^/Å^2^ (the same electron dose used in the montage tiles) and averaged for downstream .png file pre-processing and adaptive normalization to explore ultralow-dose boundary-aware flagella segmentation. This is an ideal candidate dataset for a unified model that can robustly segment thin objects in noisy cryoEM images.

#### B.2.2 Challenge Task B2: Unified Model of Continuous Flagella Segmentation in Ultralow-Dose Conditions

Flagella identification in ultralow-dose mode will be a pertinent issue in future montage tile collections. Here, we present a challenge to create a unified model for robust ultralow-dose flagella segmentation as a first step. Users can choose an ultralow-dose range for image summation from the raw cryoEM movies. Future pursuits should target bacterial flagella in a cryoEM montage dataset collection to capture the full flagella length with stitched segmentation. This additional montage stitched image set can capture the full flagella length with robust segmentation with a unified model requested above. This future dataset would thereby create synergy between noisy-image, boundary-aware models and seamless stitched segmentation routines to advance the field of automated biofilm flagella screening.

**Figure B3:**
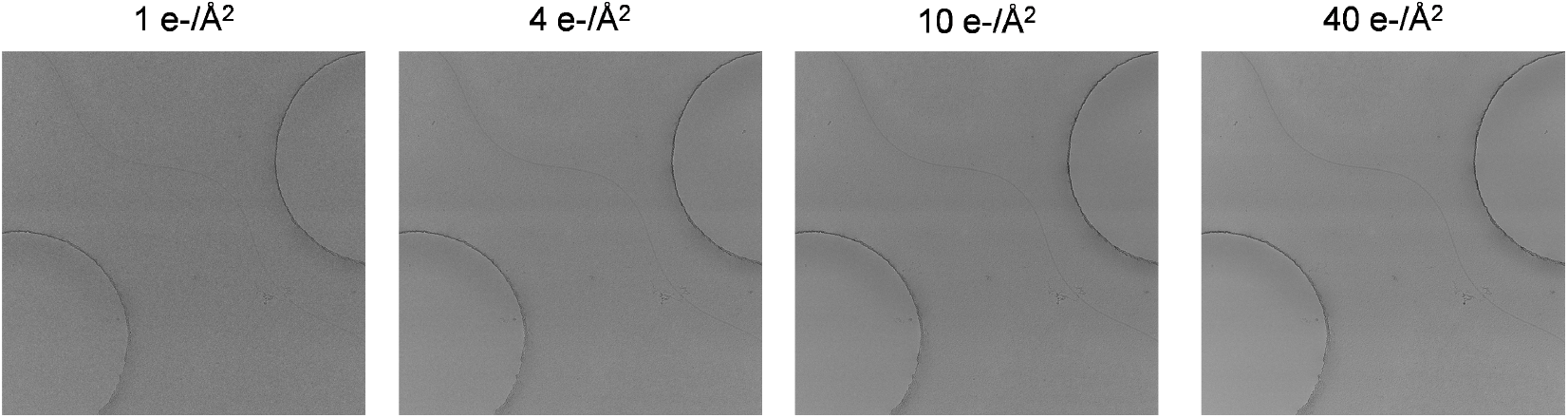
Frame partitioning for ultralow-dose with averaged cryoEM movies. From left to right, the image at 1 e^−^/Å^2^ is one frame, the 4 e^−^/Å^2^ image is four averaged frames, 10 e^−^/Å^2^ is an image of 10 averaged frames and 40 e^−^/Å^2^ is an image of 40 frames output as .png files.

**Table B1:** Summary for bacteria and flagella segmentation of datasets, including augmentations, and resizing.

| Data Structure | Classes | Electron Dose | Initial Data Size | Augmented Data Size | Augmentations and Re-sizing |
| --- | --- | --- | --- | --- | --- |
| YOLO | Flagella, bacteria | LD | 275 | 713 | Horizontal and vertical flip, crop 0% minimum and 20% maximum zoom, rotation between $-15^\circ$ and $+15^\circ$, $\pm 10^\circ$ horizontal and vertical shear, saturation between $-25\%$ and $+25\%$, $3\times$ train outputs, resized to $1024 \times 1024$ |
| COCO | Flagella, bacteria | LD | 275 | 713 | Same as above |
| YOLO | Flagella, bacteria | LD | 275 | 497 | Horizontal and vertical flip, crop 0% minimum and 20% maximum zoom, rotation between $-15^\circ$ and $+15^\circ$, $\pm 10^\circ$ horizontal and vertical shear, saturation between $-25\%$ and $+25\%$, $2\times$ train outputs, resized to $1024 \times 1024$ |
| COCO | Flagella, bacteria | LD | 275 | 497 | Same as above |

